# Archaeal chromatin ‘slinkies’ are inherently dynamic complexes with deflected DNA wrapping pathways

**DOI:** 10.1101/2020.12.08.416859

**Authors:** Samuel Bowerman, Jeff Wereszczynski, Karolin Luger

## Abstract

Eukaryotes and many archaea package their DNA with histones. While the four eukaryotic histones wrap ∼147 DNA base pairs into nucleosomes, archaeal histones form “nucleosome-like” complexes that continuously wind between 60 - 500 base pairs of DNA (“archaeasomes”), suggested by crystal contacts and analysis of cellular chromatin. Solution structures of large archaeasomes (>90 DNA base pairs) have never been directly observed. Here, we utilize molecular dynamics simulations, analytical ultracentrifugation, and cryoEM to structurally characterize the solution state of archaeasomes on longer DNA. Simulations reveal dynamics of increased accessibility without disruption of DNA-binding or tetramerization interfaces. Mg^2+^ concentration influences compaction, and cryoEM densities illustrate that DNA is wrapped in consecutive substates arranged 90^°^ out-of-plane with one another. Without ATP-dependent remodelers, archaea may leverage these inherent dynamics to balance chromatin packing and accessibility.

## Introduction

Eukaryotic genomes are orders of magnitude larger and more complex than those of archaea or bacteria. They manage their massive genomes through a hierarchical packaging scheme that utilizes histones to form nucleosomes, a complex that contains two H2A-H2B histone heterodimers flanking a central (H3-H4)_2_ heterotetramer and stably wraps ∼147 base pairs of DNA (Figure 1A) (1). While all eukaryotes invariably contain histones, genes encoding putative “minimalist” histone proteins have been identified in most archaea (2), but so far not in bacteria. While still a subject of debate, the presence of histones in the two domains of life has supported theories that eukaryotes evolved directly from an archaeon (3–9).

**Figure 1.**
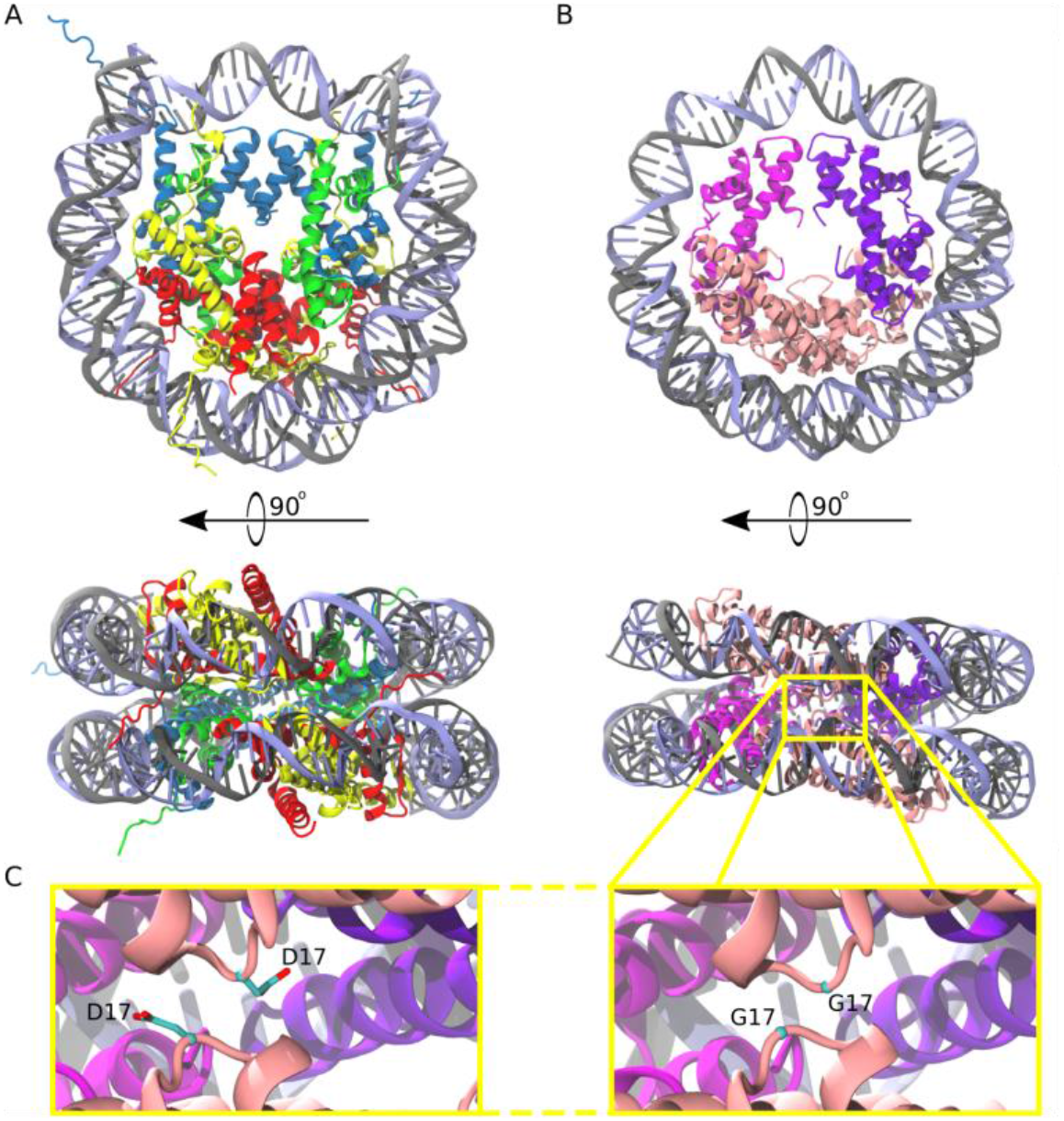
Comparison of the eukaryotic nucleosome and Arc120, an archaeasome model system containing 120 base pairs of DNA and four histone dimers. (A) The eukaryotic nucleosome (PDB 1AOI) containing two copies of H3, H4, H2A, and H2B (blue, green, gold, and red, respectively) arranged as H2A-H2B dimers flanking the (H3-H4)_2_ tetramer and binding 147 base pairs of DNA (grey and light blue). (B) The model Arc120 system, derived from the Arc90 crystal structure (PDB 5T5K), with four HMfB homodimers participating in L1-L1 stacking interactions shown in pink. (C) Enhanced views of both the wild type and G17D mutant interfaces simulated in this study.

Archaeal histones share many features with their eukaryotic equivalents, such as the three-helix histone fold motif (α1-L1-α2-L2-α3) (10), as well as obligate dimerization and transient tetramer formation (11). In both archaea and eukaryotes, histones have a preference for occupying particular sites in the genome (12, 13). Eukaryotic histones utilize highly disordered cationic N-terminal tails to regulate gene transcription and chromosome compaction via post-translational modifications (PTMs) (14–16), whereas most archaeal histones do not contain tail sequences, with the exception of some sequences found within the Asgard clade (2). Additionally, histones in eukaryotes have evolved histone fold “extensions”, unique secondary structure elements that define the outer surface of the nucleosome as well as stabilize the histone core and the final turn of nucleosomal DNA. No such diversification has yet been indentified in any of the archaeal histones.

Recently, we determined the crystal structure of *Methanothermus fervidus* histone HMfB bound to ∼90 base pairs of DNA (17, 18). HMfB is a typical representative of most archaeal histones, as it contains no tails and no histone fold extensions, and it exists as either a homodimer or heterodimer with the closely related HMfA histone (19). Nevertheless, striking similarities to the eukaryotic nucleosome were observed, including a superhelical DNA path around a histone core that is nearly identical to that in nucleosomes, promoted by conserved DNA contacts with histones, and by dimer-dimer interactions through “four-helix bundles” of α-helices. However, unlike nucleosomes, which are defined octameric particles that arrange as 10 nm “beads on a string” on long DNA fragments, crystal contacts in the archaeal system promote an extended superhelical winding of DNA around a histone core that consists of more than the canonical four histone dimers. This arrangement, which is consistent with *in vivo* footprinting results (17), places the L1 loops of every *n*^*th*^ and *(n+3)*^*th*^ dimer in close proximity to one another (Figure 1C). The importance of the L1-L1 interaction in stabilizing this arrangement in vitro and in vivo was established by substituting the highly conserved G17 residue with bulkier amino acids (e.g. G17D), resulting in transcription regulation defects.

Here, we show that archaeal histones, when assembled on long segments of DNA (> 200 base pairs), experience disparate solution dynamics compared to eukaryotic nucleosomes, and we present cryo-EM structures of both compact and open particles arranged on a 207 base pair DNA fragment. Despite many “nucleosome-like” properties, we refer to these constructs as “archaeasomes”, rather than “archaeal nucleosomes”, because of their inherent differences in higher-order structures and solution behaviors, including the ability to form histone cores with more than four dimers. We classify archaeasomes by the length of bound DNA (i.e., shorthand of “Arc120” for “archaeasome with 120 base pairs of DNA”). We use molecular dynamics (MD) simulations to show that archaeasomes can expand, like the stretching of a “slinky”, to increase overall accessibility without sacrificing histone-histone or histone-DNA interactions. Simulations also reinforce the importance of L1-L1 histone stacking interactions in regulating this dynamic equilibrium. Sedimentation velocity analytical ultracentrifugation (SV-AUC) and cryoEM highlight two distinct accessibility dynamics at play in archaeasome systems, and cryoEM analysis identifies archaeasomal states that continually wrap DNA but deflect the wrapping pathway 90^°^ out-of-plane The intrinsic dynamic behavior of archaeasome slinkies stands in contrast to eukaryotic nucleosomes, which are highly compact and require modifications and elaborate machinery to efficiently regulate chromatin accessibility, whereas archaea may utilize this inherent stochastic behavior in order to transiently access their DNA.

## Results

### Archaeasomes are highly dynamic but maintain robust dimer-dimer and dimer-DNA interactions in simulations

We applied molecular dynamics (MD) simulations to model the stability and solution dynamics of archaeasomes assembled on DNA of increasing lengths. Three system sizes were studied: an archaeasome with 90 base pairs of DNA (“Arc90”, Movie M1), one with 120 base pairs of DNA (“Arc120”, Movie M2), and one with 180 base pairs of DNA (“Arc180”, Movie M3). In *T. kodakarensis*, the G17D (G16D in HMfB) mutation in the L1 loop was previously shown to abolish the characteristic footprint of histone-based chromatin in the cell (17), and we simulated this system at the archaeasome level (“Arc120-G17D”) to elucidate the atomistic mechanisms for this observed behavior.

In each system, the DNA footprint of ∼30 base pairs per dimer was maintained on the hundreds of nanoseconds timescale. Nevertheless, the Arc90 system experienced a high degree of global dynamics. RMSD calculations of backbone atom positions showed a maximum deviation of ∼6 Å from the initial Arc90 coordinates, with trough-to-peak heights of ∼4 Å throughout the time course (Figure 2A, B). RMSF calculations identify regions of highest flexibility where the DNA binds to the terminal dimers (Figure S1), and visual inspection of Arc90 trajectories revealed that the system samples a “clamshell” motion, where terminal dimers and associated DNA fluctuate to create “closed” and “open” accessibility states (Movies M4 and M5). The equilibrium between closed and open forms was quantitated by measuring the center-of-mass distance between each DNA end and the neighboring superhelical turn of DNA, where separations of ∼16 Å and ∼30 Å are indicative of closed and open states, respectively (Figure 2C). Over the clamshell motion, the four-helix bundle interfaces that bind consecutive dimers to one another was maintained, and only small rearrangements of the local contact network were observed. Together, these data show that archaeasomes retain protein-protein contacts while still allowing for significant molecular motion.

**Figure 2.**
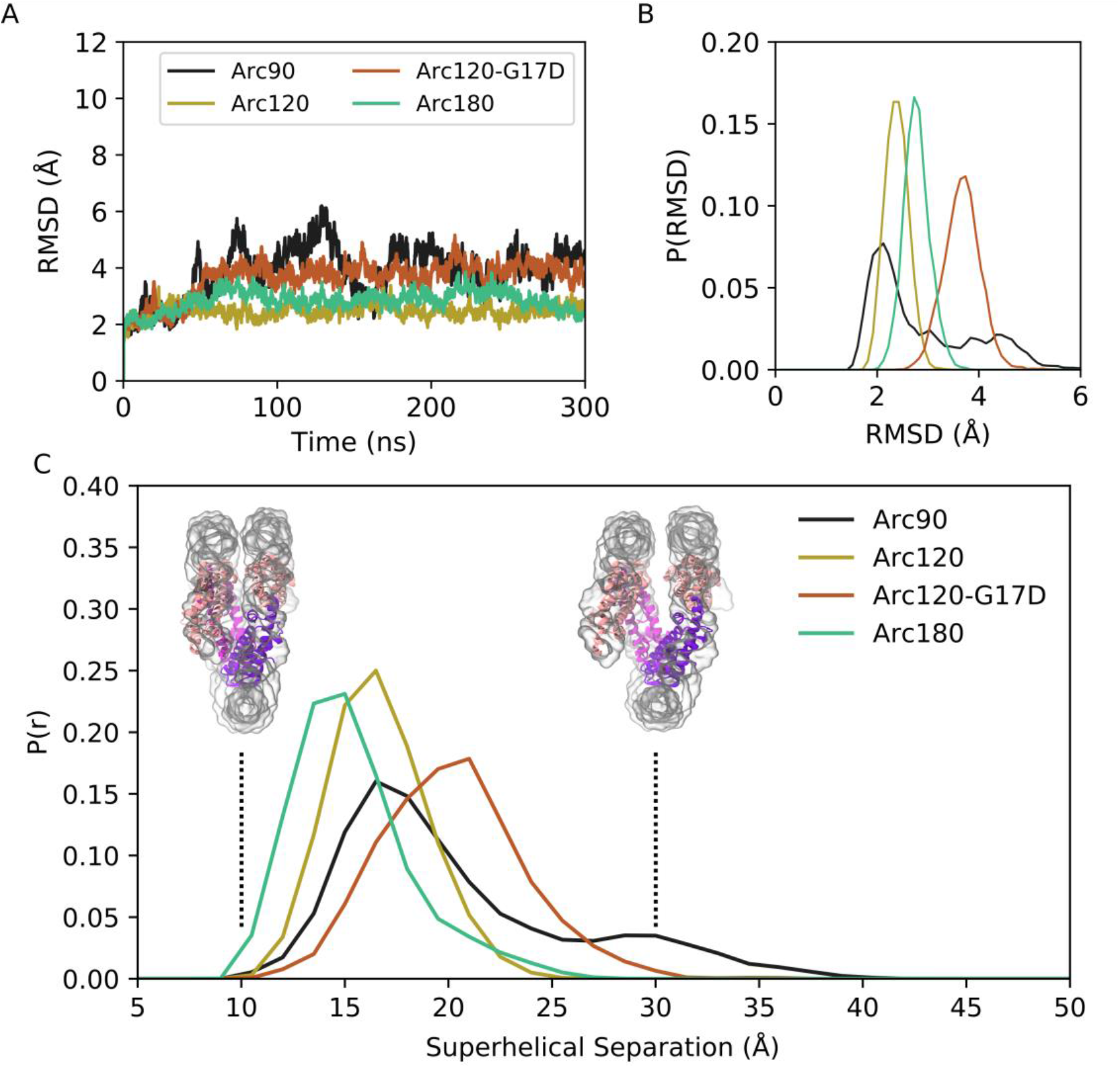
Backbone RMSD for simulated systems compared to their initial conformations. (A) Representative timeseries from each system demonstrating the largest change in structure from the initial states. The Arc90 trajectory (black) exhibits both a large peak value, as well as wide variances across the timeseries. Arc120 (gold) and Arc180 (teal) systems are significantly less dynamic, and the Arc120-G17D mutant (orange) shows an increased divergence from the initial state when compared to the wild type Arc120 system. (B) Distribution of RMSD values sampled across all three independent trajectories of each system, post-equilibration (100 ns). Arc90 displays a sampling of two different states, while each of the larger systems are unimodal. (C) Distribution curves for center-of-mass separations between DNA ends and the neighboring superhelical gyre. Representative conformations of fully closed (∼10 Å) and open (∼30 Å) states identified in the Arc120-G17D simulations are shown. Arc90 simulations (black) show a bimodal distribution between these values, but Arc120 (gold) and Arc180 (teal) systems, containing one and three L1-L1 interactions each, sample unimodal values around the closed state. The Arc120-G17D mutant (orange) samples a unimodal distribution, but indicative of a more open archaeasome than the wild type.

Simulations of Arc120 and Arc180 systems showed that L1-L1 interactions drastically reduced the population of the accessible state. Both the maximum observed RMSD and variance in the timeseries were dampened in these systems (Figure 2A, B), and RMSF calculations showed that DNA bound to the terminal dimers were less dynamic than in Arc90 trajectories (Figure S1). Similarly, the distance between DNA entry and exit sites and the neighboring DNA gyre in Arc120 and Arc180 were unimodal around closed-state values (Figure 2C), with average separations of 17.4 ± 0.2 Å and 16.1 ± 0.2 Å, respectively.

While the Arc120-G17D system has the proper number of dimers to form L1-L1 stacking interactions, the G17D mutation was designed to disfavor interactions, and this system experienced increased dynamics relative to the wild-type Arc120 trajectories. RMSD measurements showed a unimodal distribution, as was seen in the Arc120 and Arc180 simulations, but the maximum observed RMSD values were more similar to the Arc90 trajectories (Figure 2). Similarly, local fluctuations in Arc120-G17D DNA positions were increased relative to the Arc120 trajectories (Figure S1), and separation of the superhelical gyres was significantly larger than the Arc120 values (21.0 ± 0.4 Å, p-value < 0.0001). This also increased the solvent-accessible surface area by over 1,000 Å^2^ (57,316 ± 34 Å^2^ vs 58,422 ± 73 Å^2^, p-value < 0.001; Figure 3). Similar to the Arc90 system, the four-helix bundle interactions of consecutive dimers were maintained in these systems, despite the increase in DNA separation and surface accessibility

**Figure 3.**
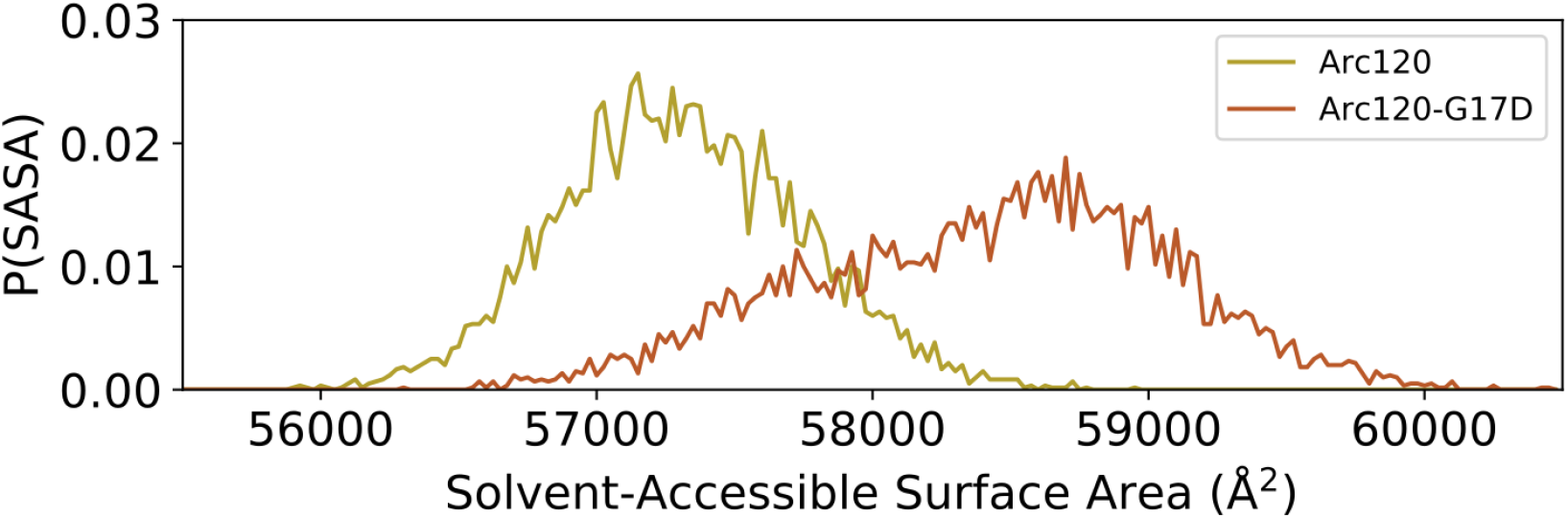
Distribution of solvent-accessible surface areas observed per frame in the Arc120 and Arc120-G17D simulations. The G17D mutation at the key L1-L1 interaction between stacking dimers significantly increases the surface area by ∼1,000 Å^2^.

Histone-DNA interactions were largely unchanged by these dynamics. Molecular Mechanics-Generalized Born Surface Area (MM-GBSA) calculations were used to estimate histone-DNA binding energies (Table 1). MM-GBSA values should not be interpreted literally, but can be used to identify qualitative and robust changes in DNA binding interactions in nucleosomes as a result of histone modifications(20–22). In agreement with previous computational studies (23), raw MM-GBSA values suggest that DNA binding strength increases as additional dimers are added to an archaeasome stack (ΔG_Arc90_ < ΔG_Arc120_ < ΔG_Arc180_). However, this may be an artifact of molecular mechanics forcefields, which are additive and thereby affected by the total number of interactions in a system. When MM-GBSA values were normalized to the number of dimers in each system, there was no net change in DNA binding ability as a function of archaeosome size. Additionally, the G17D point mutation slightly reduced the dimer-DNA interaction strength, but this difference was not statistically significant when compared to the wild-type Arc120 system (p-value of 0.157). These data, in conjunction with maintained four-helix bundle interactions, show that archaeasomes can sample open and accessible conformations without histone dissociation. Thus, the loss of the 30 base pair ladder pattern in MNase digestion of chromatin isolated from cells carrying the G17D mutation as their only source of histones may be due to increased archaeasome dynamics (Figure 3), rather than frequent histone dissociation.

**Table 1.** MM-GBSA calculations for DNA binding strength, both as raw values and normalized to the number of dimers in each system. All values are given in kcal/mol, and results are intended to be interpreted comparatively, rather than as absolute values. When normalized according to dimer count, no significant differences are observed between the wild-type histone systems of varying sizes. The G17D mutation yields a slight reduction in average calculated binding strength, but this difference is not statistically significant (p-value of 0.157). Statistical significance was calculated via t-test, and quoted error values are the standard error of the mean calculated across the three independent trajectories.

| System | $\Delta G_{\text{DNA}}$ | $\Delta G_{\text{DNA}}$ (per dimer) |
| --- | --- | --- |
| Arc90 | $-459.4 \pm 4.7$ | $-153.1 \pm 1.6$ |
| Arc120 | $-610.5 \pm 5.3$ | $-152.6 \pm 1.3$ |
| Arc120-G17D | $-580.2 \pm 15.5$ | $-145.0 \pm 6.7$ |
| Arc180 | $-906.0 \pm 5.6$ | $-151.0 \pm 0.9$ |

### Archaeasomes are inherently dynamic and accessible in solution

Our simulations provide *in silico* confirmation for the notion that archaeasomes indeed wrap several DNA turns, with more than four histone dimers, as a *bona fide* solution-state. The Arc90 and Arc120-G17D simulations suggest that archaeasomes may sample extended conformations without even partial dissociation of histones from either DNA or their histone partners. To structurally characterize these assemblies in solution, we reconstituted archaeasomes on 207 base pairs of Widom 601 DNA (Arc207) with saturating amounts of recombinant HTkA histones from *Thermococcus kodakarensis*, which are very similar to HMfB histones (overall 59% identity, 81% similarity, with higher conservation in sequences involved in histone-histone and histone-DNA interactions) (17). The solution behavior of the Arc207 construct was then analyzed by single particle cryoEM and sedimentation velocity analytical ultracentrifugation (SV-AUC). As controls, we also collected SV-AUC traces from eukaryotic (*Xenopus laevis*) nucleosomes reconstituted on 147 base pairs of Widom 601 DNA.

Gel shift assays showed that full complex saturation occurs when DNA and histones are mixed at the previously reported stoichiometric limit (∼30 bp per dimer; Figure S3). SV-AUC traces show that the Arc207 complex sediments homogeneously at ∼10.6 S (Figure 4, circles), and the Nuc147 control sediments more rapidly than the Arc207 system at ∼11.2 S (Figure 4, squares). As sedimentation is dependent on both molar mass (higher mass yields faster sedimentation) and shape anisotropy (higher anisotropy yields increased drag and slower sedimentation), the relatively lower sedimentation rate of the Arc207 sample has one of two interpretations: either its mass is less than that of a nucleosome, or the particle is more extended in solution, exhibiting higher shape anisotropy and drag.

**Figure 4.**
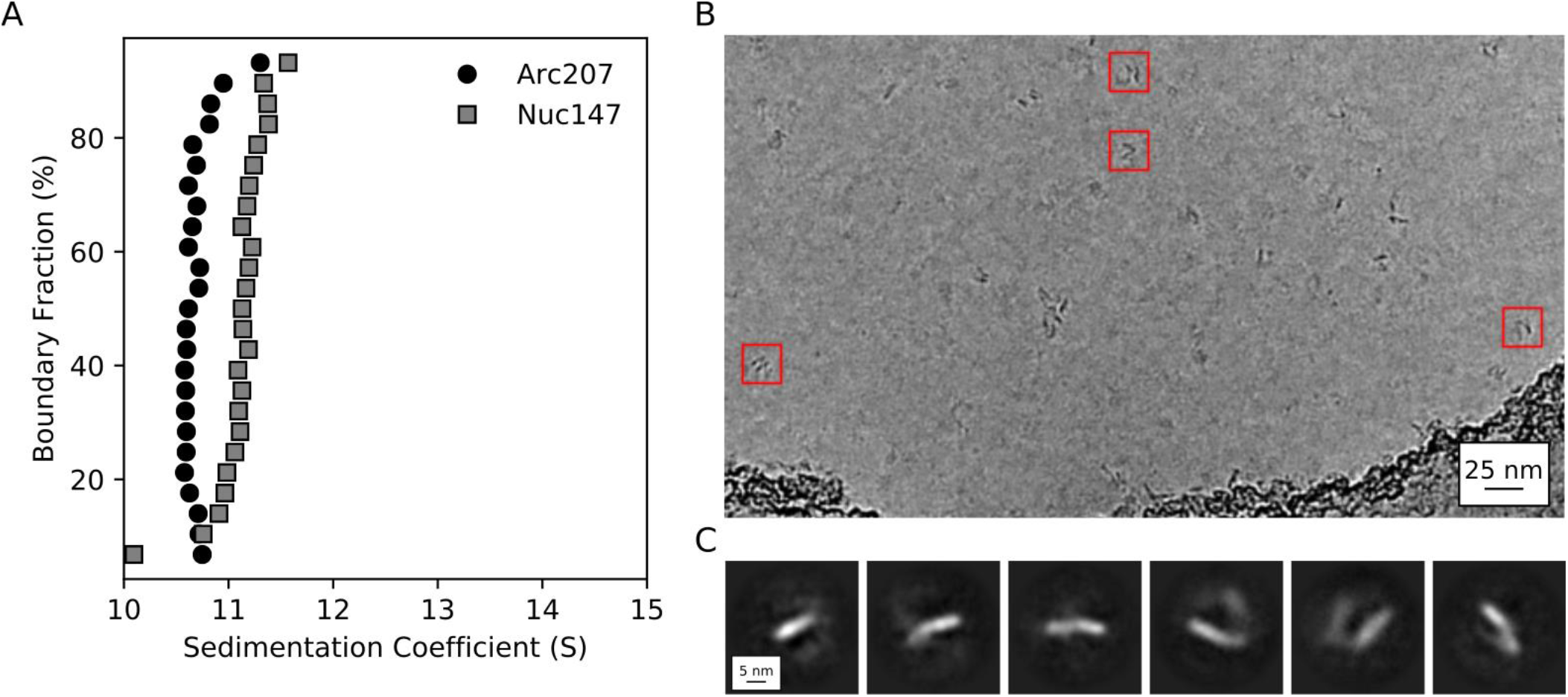
Biophysical characterization of archaeasomes in vitro. (A) van Holde-Weischet plots of Arc207 (circles) and Nuc147 (squares) samples. While Arc207 has a larger mass than the eukaryotic Nuc147, it sediments more slowly, indicative of increased drag caused by extended particle configurations. (B) Representative de-noised cryoEM micrograph of Arc207 sample. Particles are observed with DNA wrapped in a nucleosome-like pattern, but a significant separation between neighboring DNA turns is observed. (C) Two-dimensional classifications of particles extracted from these micrographs. No tightly wrapped DNA is identified, in agreement with our SV-AUC inferences of an extended Arc207 particle shape in comparison to compact Nuc147.

The mass and anisotropy of each molecule were modeled from SV-AUC traces using the Monte-Carlo-coupled Genetic Algorithm (GA-MC) module of Ultrascan III (Table 2). The molecular weight for both particles was slightly overestimated, consistent with a systemic error in specific volume estimation for protein-DNA complexes, which can propagate to modest errors in molecular weight estimation (24). Nevertheless, these data show that the Arc207 complex is fully saturated with histones, that the total mass is consistent with seven histone dimers assembled on 207 bp of DNA, and that its mass is greater than that of Nuc147. This suggests that the Arc207 system is more anisotropic than Nuc147 in order to satisfy the decrease in sedimentation rate, which is confirmed by the frictional ratios (f/f_o_) estimated from GA-MC calculations (1.94 and 1.55 for Arc207 and Nuc147, respectively). For reference, the f/f_o_ value of free 207 DNA is 3.14 so the Arc207 complex is more compact than free DNA but not as compact as the eukaryotic nucleosome.

**Table 2.** SV-AUC properties of Arc207 and Nuc147 molecules. Despite the higher mass of the Arc207 complex, it sediments slower than the Nuc147 system due to increased particle elongation (frictional ratio, “f/f_o_”). “MW_the_” denotes the theoretical molecular weight, and “MW_GA-MC_” corresponds to the molecular weight derived via GA-MC analysis. Parenthetical values outline the 95% confidence interval of each GA-MC value.

|  | Nuc147 | Arc207 | 207 DNA |
| --- | --- | --- | --- |
| $MW_{the}$ (kDa) | 200.3 | 231.4 | 127.8 |
| $MW_{GA-MC}$ (kDa) | 218.6<br>(207.2, 229.9) | 266.5<br>(256.9, 276.1) | 156.6<br>(152.4, 160.1) |
| $f/f_0$ | 1.55<br>(1.49, 1.61) | 1.94<br>(1.89, 1.99) | 3.14<br>(3.08, 3.20) |
| Sedimentation Coefficient (S) | 11.2 | 10.6 | 6.1 |

We next utilized single particle cryoEM to capture the three-dimensional structure of the Arc207 complex. Our simulations of the similarly sized Arc180 system predicted that the complex would be compact, but SV-AUC modeling conversely suggests that Arc207 forms an extended conformation. CryoEM data shows that the Arc207 system is indeed open, as it exhibits nucleosome-like particle dimensions with increased spacing between neighboring DNA gyres (Figure 4B), in agreement with SV-AUC data. This is also apparent in the two-dimensional classifications (Figure 4C). Because of the apparent structural heterogeneity of this assembly, we were unable to extract three-dimensional conformations from the dataset, even at low resolution.

### Archaeasomes compact but do not oligomerize in the presence of Mg^2+^

Eukaryotic chromatin fibers can be compacted *in vitro* through the addition of divalent cations such as Mg^2+^ (25). To test whether this also applies to archaeal chromatin, we analyzed Arc207 samples in the presence of 0, 1, 2, 5, 7, 8, and 10 mM MgCl_2_ by SV-AUC, and we observed changes in both sedimentation coefficient and frictional ratio (Figure 5). Arc207 samples exhibit an increase in sedimentation rate with increased Mg^2+^ concentration from 0 to 5 mM, with no additional increase between 5 to 10 mM MgCl_2_ (Figure 5A, top). In comparison, the nucleosome system shows little to no change in sedimentation from 0 to 2 mM MgCl_2_ (Figure 5A, bottom), and start to self-associate (aggregate) at concentrations above 2 mM Mg^2+^.

**Figure 5.**
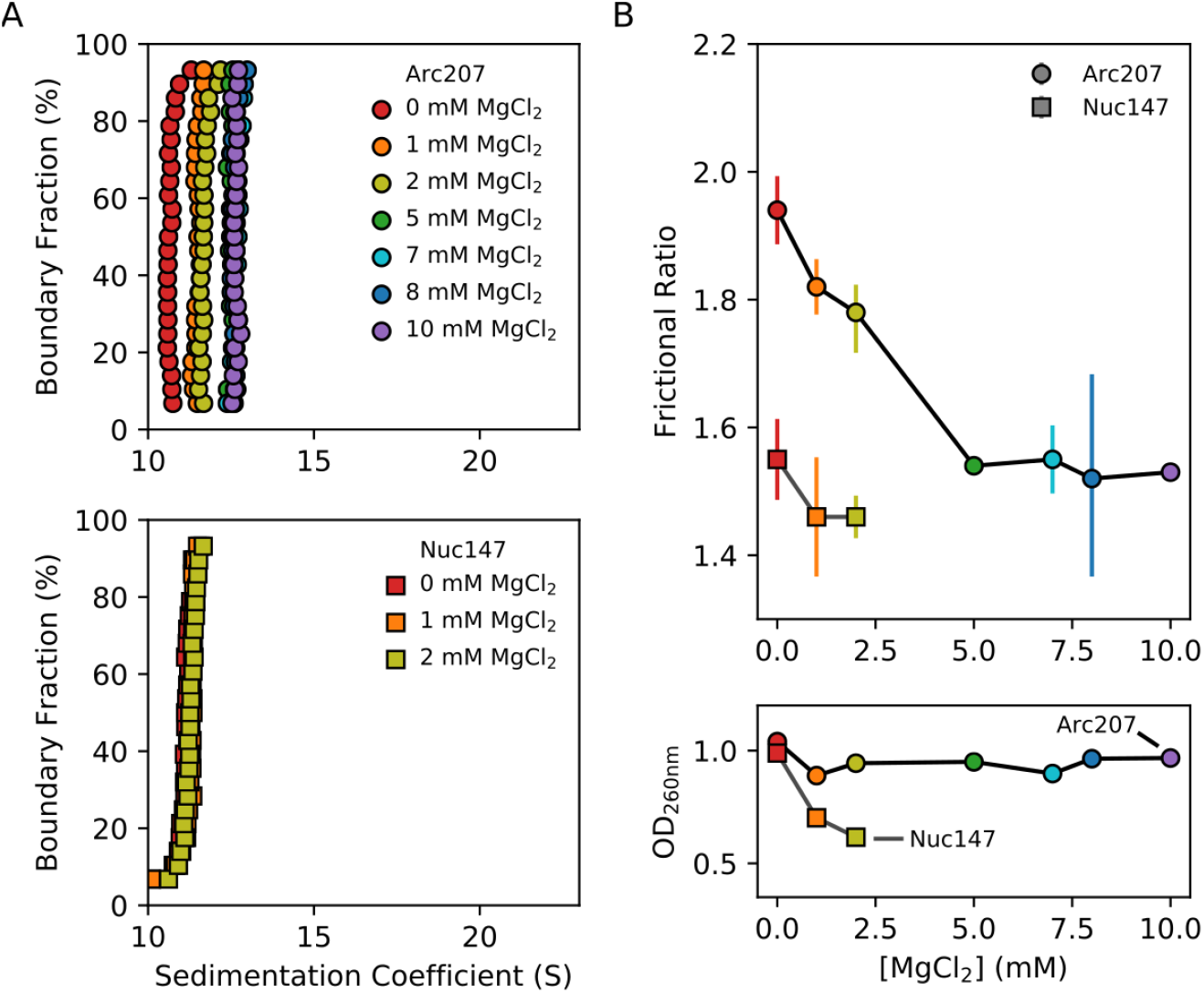
Effects of Mg^2+^ concentration on sedimentation behavior of Arc207 and Nuc147 samples. (A) van Holde-Weischet plots of (top) Arc207 and (bottom) Nuc147 samples. Increases from 0 to 5 mM MgCl_2_ results in an increase in Arc207 sedimentation rate, but little change in Nuc147 sedimentation. (B) Effects of MgCl_2_ concentration on (top) frictional ratio and (bottom) absorbance at 260 nm for Arc207 (circles) and Nuc147 (squares). Frictional ratios of Arc207 samples follow the same profile as sedimentation rate, where samples compact from 0 to 5 mM MgCl_2_ but with little change from 5 to 10 mM MgCl_2_. Similarly, Nuc147 samples showed very modest changes in compaction. OD_260nm_ measurements show Nuc147 particles aggregating as a result of increased MgCl_2_ concentration, but Arc207 samples compact without losses to aggregation.

GA-MC analysis of the Arc207 traces shows the same correlation between f/f_o_ and MgCl_2_ concentration that is observed for the sedimentation rate, with apparent compaction from 1 to 5 mM MgCl_2_ and no further reduction beyond 5 mM MgCl_2_ (Figure 5B, top). A modest reduction in f/f_o_ is observed for the Nuc147 sample, but the most notable differences between the archaeasome and nucleosome systems is the loss of nucleosome sample due to aggregation. For Nuc147, OD_260_ values rapidly decline with increased Mg^2+^, and no appreciable sample remains at 5 mM MgCl_2_ and above (Figure 5B, bottom). In contrast, the Arc207 complex displays no significant losses, even at 10 mM MgCl_2_. This shows that, unlike eukaryotic chromatin, divalent cations compact archaeasomes without promoting fiber-fiber association.

### Archaeasomes exist in open and closed states

Single particle cryoEM was utilized to determine the three-dimensional structure of archaeasomes in the presence of 5 mM MgCl_2_. Particles appeared more defined than the samples without MgCl_2_, and individual particles with fully wrapped DNA conformations are distinguishable from other states where density is visible as perpendicular extensions to the wrapping plane (Figure 6A, blue and gold boxes). A total of 1,879,294 particles were identified and classified according to a neural network trained on manual particle selections. The two-dimensional classes of this dataset confirm the presence of open and closed forms of Arc207 (Figure 6C, D). We separated these particles into independent 3D classes and refinement schemes, which yielded 80,609 particles in the compact form and 5,959 particles in the open form. Many more particles could have been included in each class, especially the open form, but we were conservative to ensure that no interactions with neighboring particles affected their configuration. Refinement of these densities yielded maps at 9.5 Å and 11.5 Å resolution for the closed and open forms, respectively (Figure 6E, F).

**Figure 6.**
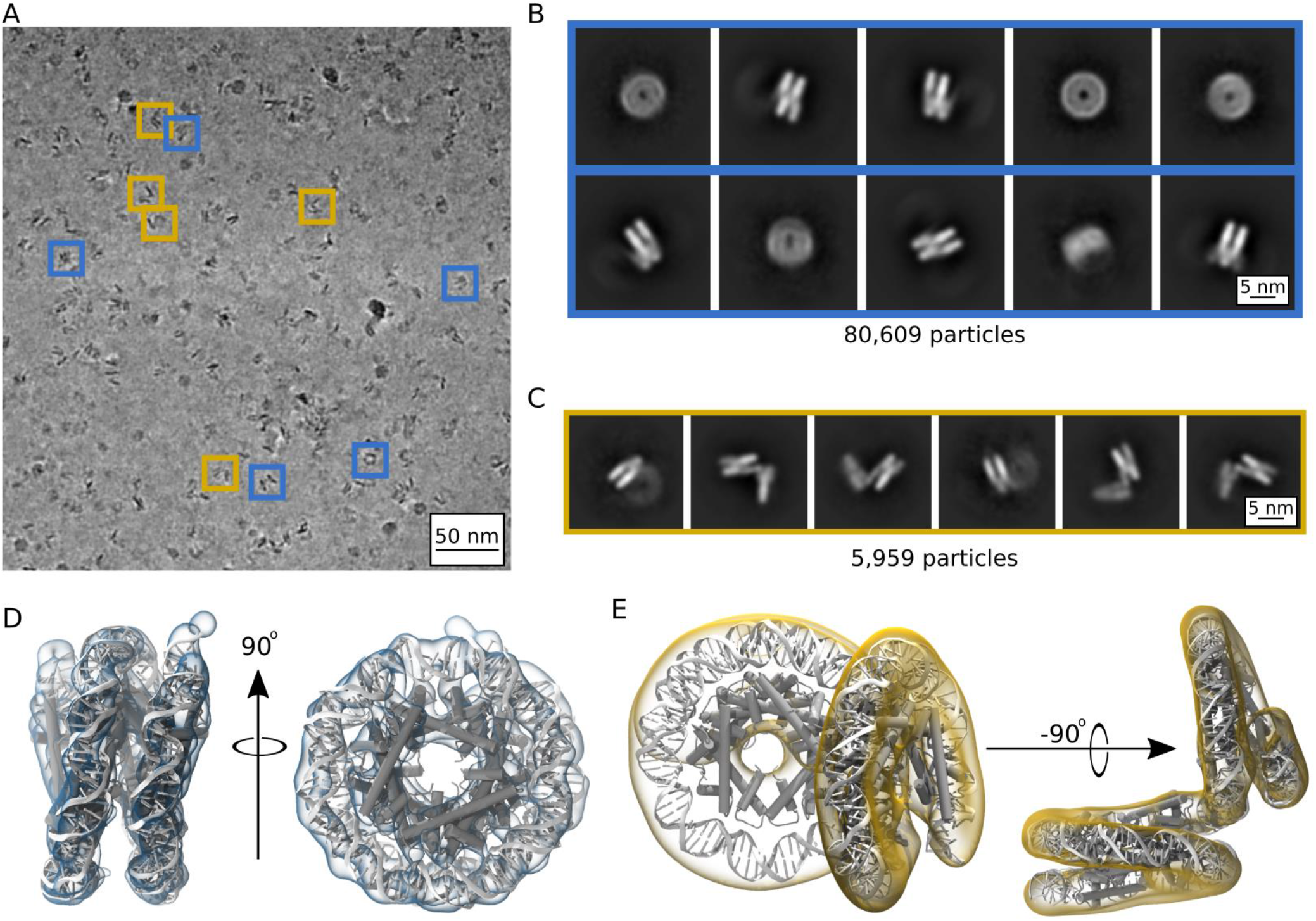
Single particle cryoEM analysis of the Arc207 complex in the presence of 5 mM MgCl_2_. (A) Representative micrograph, denoised for visibility. Particles are observed with both compact wrappings of the 207 base pair DNA fragment (blue boxes) as well as particles showing out-of-plane “lid” extensions of DNA from similarly wrapped “cores” (gold boxes). Also shown are two-dimensional classes derived for the (B) closed and (C) open particle classifications. Three-dimensional densities from these classes are shown in (D) and (E), respectively. The closed-form particle was fit by rigid-body docking Arc150 coordinates extracted from our Arc180 simulations, and the open-form particle was fit by rigid-body docking separate Arc90 and Arc120 components and bridging the connecting DNA through energy minimization.

At 9.5 Å resolution, secondary structure and side chain assignment is not possible for the closed form structure. However, key features of the archaeasome previously seen in the crystal structure are discernible from the density, such as DNA wrapping around a core of five histone dimers and periodic, overlapping densities that we associate with α2, the longest of the core α-helices in each histone (Movie M6). Even though Arc207 (207 base pairs of DNA bound to 7 histone dimers) were deposited on the grid, our refined density of the closed state describes only 150 base pairs of DNA and 5 histone dimers (Movie M7). As such, we trimmed the final frame from one of our MD simulations of the Arc180 complex down to an Arc150 complex and docked it to the map using the “fit in map” function of Chimera. We simulated density at 8 Å resolution for the MD-derived structure, which yields a correlation coefficient of 0.85 with the empirical map. This shows that our MD simulations describe a compact archaeasome with high fidelity.

The open form density has sufficient volume to be fit with the full 207 base pair DNA and 7 histone dimers (Movie M8). The larger portion of the volume is best described by four histone dimers binding ∼120 base pairs of DNA in a nucleosome-like structure, while the smaller portion can be fit with three histone dimers binding ∼90 base pairs. While the connecting density suggests only moderate bends in the helical axis, the wrapping pathways of the sub-units result in the two faces being arranged at a near 90° angle. The continuity of the DNA between the two moieties was confirmed by modulating the electron density contours (Movie M9). Structural models were generated by rigid-body docking separate Arc120 and Arc90 subunits into the associated density, and the connecting DNA was energy minimized and subjected to a short MD simulation to confirm that the connecting DNA segment is reconcilable with B-form DNA parameters (Figure S4).

To confirm that the structures described by the EM densities also exist in solution, and to correlate these findings with our SV-AUC analyses of particle compaction, we utilized the SoMo plugin of UltraScan to calculate sedimentation properties from our derived models. For the open-form Arc207 model, with the Arc90 “lid” bending ∼90^°^ from the Arc120 “core”, we calculate a frictional ratio of 1.53, in close agreement with the 1.55 value extracted from the experimental traces. For the closed form, we assumed that the remaining 60 base pairs of DNA (and 2 histone dimers) extend from the observed density, but are too disordered to be seen in the class averages. Close inspection of the “closed state” two-dimensional classes indeed shows additional out-of-plane density that is consistent with the missing two dimers and ∼60 base pairs of DNA, albeit at low contrast relative to background (Figure 7C). To model this particle, we extended the closed-form model to contain an Arc60 subunit similarly positioned 90^°^ out-of-plane with the resolved Arc150 density and again find a similar value of 1.50. In contrast, models with continual wrapping (as observed in the crystal structure) yield a frictional coefficient of 1.36. These data together show that archaeasomes on an extended DNA fragment can be viewed as a distribution of archaeasome subunits that can open up to a ∼90^°^ angle.

## Discussion

Using *in silico* and *in vitro* approaches, we investigated the structure and dynamics of the ‘slinky-like’ architecture of histone-based archaeal chromatin. SV-AUC and cryoEM both show that the archaeasome, containing more than the characteristic four histone fold dimers observed in nucleosomes, is inherently dynamic and exists in open and closed states while maintaining DNA-histone interactions. Archaeasome compaction can be stabilized with divalent cations. In absence of archaeal ATP-dependent chromatin remodeling factors (large machines that regulate chromatin access in eukaryotes), this architecture provides an alternative mechanism for compacting chromatin and adjusting genome accessibility.

Our cryoEM structures show that archaeosomes are metastable in the absence of divalent cations. Even in the presence of 5 mM Mg^2+^, archaeasomes exhibit closed and open configurations. Indeed, our structures contain only four to five histone dimers arranged in a continuous helical ramp. Beyond that, the helical histone ramp is disrupted at the four-helix bundle interface by an outward rotation of the remaining two to three histone dimers and attached DNA. The existence of these structures in solution is supported by analysis of the hydrodynamic parameters of archaeosomes obtained from SV-AUC. Extending this arrangement to even longer DNA, one could picture particles that wrap anywhere from ∼90 to ∼150 base pairs (using three to five histone fold dimers) arranged at ∼90 degree angles with respect to each other, connected by minimal linker DNA. This arrangement may leave the major groove of the connecting DNA susceptible to nuclear factors, such as micrococcal nuclease or transcription factors, and explains the ∼30 bp MNase digestion ladder observed in native archaeal chromatin.

Consistent with this interpretation, molecular dynamics simulations of Arc90 assemblies (with three histone fold dimers) displayed dynamic breathing at terminal sites of the complex, while histone-histone and histone-DNA contacts are maintained. Models constructed from repeats of this conformation agree with particle conformations extracted from SV-AUC hydrodynamic parameters, as well as direct images and two-dimensional class averages from cryoEM data gathered in the absence of Mg^2+^. There, multiple turns of DNA, fully saturated with histones, were observed but with considerable flexibility and distance between neighboring gyres. In contrast, Arc120 and Arc180 simulations (containing four and five histone dimers, respectively) did not exhibit this degree of dynamic behavior. However, these simulations may have been artificially biased for the closed configuration, because the starting structures had intact L1-L1 interactions. Separation of these contacts may require more than several hundred nanoseconds of computational sampling, and simulations of the Arc120-G17D system indeed show appreciable separation between DNA turns by weakening this interface while maintaining four-helix bundle interactions.

In eukaryotes, chromatin structure can be further modulated by the incorporation of histone variants (26). While in vivo studies have identified disparate roles of variant histones in archaea (27), very little is known about the structural modifications that they invoke. In a recent study by Stevens et al. (28), molecular dynamics trajectories showed that substituting major type histones in *Methanosphaera stadtmanae* with a “capstone” paralog, a histone variant with sequence modifications at the four-helix bundle, disrupts the tetramer interface and separates histone dimers on a continuous DNA fragment, thereby limiting archaeasome size. In the context of the capstone model, our EM-derived structures suggest that destabilizing the four-helix bundle interface may yield a subtle increase in DNA accessibility through increased exposure of the major groove, and without the need to fully displace the archaeasome subunit. As we have shown, this dynamic behavior is already sampled by major-type archaeal histones deposited on a continuous DNA fragment, and substitution of terminal histones with capstone-containing dimers will further bias the assembly towards the open configuration. In this way, lengths of DNA that would typically contain a single archaeasome unit of considerable length would instead be segmented in to several sub-archaeasomes configured in repeat perpendicular arrangements that modestly expose bridging DNA segments. Additionally, high rigidity in certain DNA sequences may inherently poise these segments as “linker” sequences, and the frequency of these less flexible regions would similarly influence the number of 90^°^ - oriented subunits present along the genome.

Our SV-AUC traces show that Arc207 assemblies exhibit no signs of self-association upon the addition of up to 10 mM MgCl_2_, unlike eukaryotic nucleosomes and nucleosome arrays, where this favors extensive inter-nucleosome and inter-array interactions and aggregation (25). Instead, these conditions bias individual archaeasomes toward more compact states without promoting inter-archaeasome interactions. Interestingly, estimates of Mg^2+^ concentration within *T. kodakarensis* cells have been quoted at ∼120 mM (29), much higher than measurements of Mg^2+^ in mammalian nuclei (16-18 mM) (30). Per dimer, archaeal histones provide weaker electrostatic screening of DNA than eukaryotic histones, due to their relatively lower isoelectric points (pI of ∼8 vs ∼11), and the greater overall negative charge of archaeasomes may encourage rapid sequestering of any free Mg^2+^ ions in the cell. This would induce a gradient-based mechanism for rapid and excessive uptake of cations to prevent chromatin compaction from monopolizing the Mg^2+^ necessary for enzyme action. On the other hand, the ability of *T. kodakarensis* archaeasomes to compact but not aggregate as a result of elevated Mg^2+^ concentration may be a direct result of evolving in high-salt environments, as regulating between internal and external concentrations may have been more energetically demanding than simply adapting their chromatin structure. More specific measures of local Mg^2+^ concentration in archaeal chromatin may help differentiate these two mechanisms.

Archaeal histones were successfully expressed in *E. coli* cells and were found to coat bacterial DNA (23). Surprisingly, this resulted in only minor perturbations of cell growth, despite *E. coli* lacking the evolutionary machinery to handle histones. Furthermore, *in vitro* assays have shown that transcription through archaeasomes is slowed but not altogether stopped (31). In eukaryotes, navigation of polymerases through chroimatin is assisted by histone chaperones and ATP-dependent remodeling factors (32–34), but no known homologs to these complexes have been identified in archaea. The inherent dynamics of archaeasome-based chromatin may thereby limit, and not entirely prevent, access to chromatin by the sporadic appearance of near-zero linker DNA and Arc60 or Arc90 substates. The degree to which DNA rigidity, variant histones, or unknown histone- and DNA-binding proteins regulate further accessibility is an exciting future of research.

## Methods

### Molecular Dynamics Simulations

Archaeasome systems of varying sizes were constructed from the crystal structure containing ∼90 bp of DNA bound to histone HMfB (PDB 5T5K). Systems containing ∼90 base pairs of DNA and three histone dimers (Arc90), ∼120 base pairs of DNA and four histone dimers (Arc120), and ∼180 base pairs of DNA and six histone dimers (Arc180) were created as prescribed by the crystal lattice. The Arc90 system is intended to represent the “fundamental unit” of archaeasome-based chromatin, as it is the crystallographic unit of the solved structure as well as the smallest DNA protection footprint observed by MNase digestion (17). Additionally, an Arc120 system with the G17D mutation was also generated (Arc120-G17D), in analogy to the destabilizing G17D mutation that was previously studied *in vivo*. The overhanging DNA bases that formed crystallographic contacts were removed in our simulations. Each system was neutralized and solvated in a TIP3P box of 100 mM NaCl (35), and masses were repartitioned from heavy atoms to covalently bonded hydrogen atoms to allow for the use of a 4 fs timestep (36). Parameters for protein atoms were taken from the Amber FF14SB forcefield (37), DNA parameters were taken from the Amber bsc1 forcefield (38), and ions were parameterized according to the modifications of Young and Cheatham (39).

Systems were then energy minimized for 5,000 steps while constraining solute heavy atoms with a 10 kcal/mol/Å^2^ harmonic potential, followed by 5,000 steps without restraints. After energy minimization, three independent simulations of each system were conducted according to the following protocol: simulations were heated from 10 K to 300 K over the course of 50 ps in the NVT ensemble with heavy atom restraints applied, system densities were equilibrated and positional restraints were slowly released over the course of 200 ps in the NPT ensemble (target pressure of 1 atm), and simulations were extended for 300 ns without positional restraints in the NPT ensemble. During these production simulations, terminal base pair fraying was removed from the simulation by reinforcing hydrogen bonding in the terminal base pairs. No restraints were applied when participating atoms were within 3.5 Å of one another, but a harmonic potential was applied when participating atoms spread beyond this cutoff (force constant = 5.0 kcal/mol/Å^2^). Minimization and MD simulations were conducted in the pmemd engine (v18), with CUDA acceleration utilized for the simulations (40).

### Simulation Analysis

System equilibration was monitored through RMSD calculations of backbone atom positions, and local flexibility was measured via the root mean-squared fluctuation (RMSF) of the backbone. DNA breathing dynamics were quantified using the center of mass distance between the terminal base pairs and the neighboring superhelical turn. Complex stabilities were assessed using the MM-GBSA method with the igb5 solvent parameters and mbondi2 atomic radii (41). Trajectories and single frames were rendered using VMD, and structural analyses (RMSF, RMSD, DNA breathing) were calculated using cpptraj. Statistical significance between populations were determined by unpaired t-test.

### In Vitro Sample Preparation

HTkA histones were provided by the Histone Source (CSU Fort Collins, CO), and the 207 base pair, Widom 601-derived sequence was purified as described (17, 42). Archaeasome complexes (Arc207) were formed by mixing 207 bp DNA with HTkA histones at a 1:7 molar ratio of DNA to histone dimer in a buffer containing 100 mM KCl and 50 mM Tris (pH 8.0) and incubated for 20 minutes at room temperature. Samples were then dialyzed in buffers containing 0, 2, 5, 7, or 10 mM MgCl_2_ at a volume ratio of 1:1000 two times, first for two hours and then overnight, as preparation for subsequent measurements. As histone stocks are suspended in high glycerol, the dialysis process simultaneously served to effectively remove glycerol.

### SV-AUC

Sedimentation velocity analytical ultracentifugation (SV-AUC) measurements were conducted in absorbance mode (λ = 260 nm). Samples were loaded in to an An60Ti rotor in 400 µL cells with 2-channel Epon centerpieces and then spun at 35,000 rpm at 20 °C in a Beckman XL-A ultracentrifuge. Partial specific volumes of the samples were determined using UltraScan3 (v4.0)(43). Time and radially invariant noises were subtracted through two-dimensional sediment analysis (2DSA), and the final 2DSA model parameters were used to initialize genetic algorithm and Monte Carlo analyses. Sedimentation coefficients were determined using van Holde-Weischet analysis (in Svedberg units, corrected to solvent conditions of water at 20 °C), and molecular weights and frictional ratios (f/f_o_ - the degree of elongation/flexibility in comparison to ideal spherical particles) were determined from genetic algorithm analysis of SV-AUC traces. Theoretical f/f_o_ values for modeled structures were derived using the SoMo plugin of UltraScan, where bead models were created through the “SoMo Overlap” scheme and hydrodynamics were calculated with the ZENO algorithm (44, 45).

### CryoEM Grid Preparation and Collection

After overnight dialysis (100 mM KCl, 50 mM Tris pH 8, 5 mM MgCl_2_), Arc207 samples were concentrated to 0.9 mg of DNA per mL, and 4 µL of sample was deposited onto glow-discharged grids. Samples were screened on copper Cflat (1.2/1.3) grids, and formaldehyde-cleaned Quantifoil (2/2) grids were used for final data collection in order to increase image acquisition speed through a “2×2” collection scheme, where a single defocus level is used for a cluster of four grid holes. Each grid was manually plunge frozen in liquid ethane and stored in liquid nitrogen. Datasets were collected on a FEI Tecnai F20 equipped with a Gatan K3 camera at 29,000x magnification in non-super resolution mode (yielding 1.291 Å per pixel), 200 kV accelerating voltage, dosage rate of ∼1 e^-^/ Å^2^, and 50 frames per micrograph stack. A total of 5,388 image stacks were collected for subsequent three-dimensional analysis. An Elsa Cryo-Transfer Holder (Gatan, Inc.) and defocus range of −1.2 to −2.6 μm was used.

### Single Particle Analysis of CryoEM Data

Using the Relion interface (v3.0) (46), micrographs were motion corrected using MotionCorr2 (47), and CTF parameters were estimated from gCTF (48). Ten micrographs were then randomly selected from a collection of motion-corrected micrographs and denoised using the janni_denoise.py function of the SPHIRE-crYOLO particle-picking pipeline (49). Particles were manually picked from these denoised micrographs and used to train a neural network through crYOLO for picking particle coordinates across all micrographs, where the final particle predictions utilized the same denoising process (50).

Densities shown here are the result of refinements to the largest dataset, collected in the 2×2 scheme. In total, 1,879,294 particles were predicted by crYOLO and extracted from the motion-corrected (non-denoised) micrographs using Relion and imported to CryoSparc (v2.12.4) for two-dimesional (2D) classification (51). 2D classes were manually filtered to remove particles containing primarily noise or interactions with overlapping neighbor particles. Subsequent 2D classifications showed two predominant particle types: a “closed” archaeasome state characterized by tightly wound DNA, and an “open” archaeasome state showing nucleosome-like DNA arrangements with distinct out-of-plane densities. These classes were then separated from one another for three-dimensional (3D) classification and refinement by CryoSparc. The closed-state density was refined through Bayesian Polishing of the contributing particles in Relion, but the open-state density saw no benefit from polishing and the CryoSparc-derived density is reported. Figures and movies were generated with VMD (v1.9.3) (52).

### Modeling of EM Densities

The closed-form density (containing volume of five histone dimers and ∼150 bp of DNA) was modeled by extracting five dimers and ∼150 bp of DNA from the end state of a randomly selected Arc180 simulation. Simulation coordinates were fit to the EM-derived density through rigid body docking in Chimera. The open-form density was modeled by first docking Arc90 and Arc117 constructs in the smaller and larger volumes, respectively. Then, the bridging DNA segments were ligated via tleap and energy-minimized via pmemd in implicit solvent (Amber FF14SB protein forcefield, DNA bsc1 parameters, and igb5 implicit solvent model with mbondi2 atomic radii modifications and a 100 mM monovalent salt environment). Calculation of nucleic acid geometry parameters was conducted with cpptraj.

## Acknowledgements

We thank Garrett Edwards for valuable conversations regarding SV-AUC experiments and analyses, as well as Pamela Dyer and Uma Muthurajan for discussions about sample preparation. We also thank Alison White for purified DNA stocks, and Kathryn M. Stevens and Tobias Warnecke for constructive feedback on this work.

Work in the Weresczyznski group was supported by an NSF Career Award (1552743) and an NIH NIGMS R35 award (R35GM119647), and funding for the Luger lab is provided by the Howard Hughes Medical Institute (HHMI). SB was funded in part by both HHMI and the NIGMS R35 Award to JW, as well as by an NIH NIGMS NRSA fellowship (F32GM137496). The contents of this article are the sole responsibility of the authors and do not necessarily represent the official views of the National Institutes of Health.

MD simulations were conducted, in part, on an SDSC Comet GPU allocation to SB through the XSEDE environment, which is supported by NSF grant ACI-1548562 (53). Electron microscopy was done at the University of Colorado, Boulder, EM Services Core Facility in the MCDB Department, with the technical assistance of facility staff. Calculations for modeling SV-AUC data were performed on the UltraScan LIMS cluster at the Bioinformatics Core Facility at the University of Texas Health Science Center at San Antonio and multiple high-performance computing clusters supported by NSF XSEDE Grant #MCB070038 (to Borries Demeler).

## Author Contributions

Samuel Bowerman – Conceptualization; Methodology; Validation; Formal Analysis; Investigation; Data Curation; Writing – original draft; Writing – Review & Editing; Visualization; Resources; Funding Acquisition

Jeff Wereszczynski – Conceptualization; Writing – original draft; writing – review & editing; Resources; Funding Acquisition; Supervision

Karolin Luger – Conceptualization; Writing – original draft; writing – review & editing; Resources; Funding Acquisition; Supervision

## Movie Captions

**Movie M1. Rotating view of the Arc90 system**. This system contains 3 histone dimers and no stacking interactions.

**Movie M2. Rotating view of the Arc120 system**. This system contains 4 histone dimers and one L1-L1 stacking interaction. Both the wild-type and G17D mutant of this system were simulated.

**Movie M3. Rotating view of the Arc180 system**. This system contains 6 histone dimers and three L1-L1 stacking interactions.

**Movie M4. Face-view of dynamics observed in a single Arc90 trajectory**. While the terminal dimers exhibit large fluctuations around the central histones, four-helix bundle interactions are maintained.

**Movie M5. Side-view of dynamics observed in a single Arc90 trajectory**. The dynamics of the terminal dimers creates “open” and “closed” states in a “clamshell-like” motion.

**Movie M6. Derived EM density for the closed-form state of the Arc207 complex**. By varying the contour levels of the density, we find that the strongest contributors to the EM volume are around the outside of the molecule (attributed to DNA), as well as overlapping periodic densities in the histone core (attributed to each histone’s α2 helix).

**Movie M7. EM density of closed-form state of Arc207 complex (clear blue) docked with Arc150 complex (grey) extracted from MD simulations of Arc180 complex**. Docking of the Arc150 construct shows that our closed-form density cannot capture the full complex deposited on the grid but only ∼150 base pairs of DNA and 5 histone dimers. Additional density can be seen to extend along the DNA path beyond the bound core, suggesting a possible continuation of linker DNA that could extend to the potential tetramers observed in the 2D classes (see main text for details).

**Movie M8. EM density of open-form state of Arc207 complex (clear gold) docked with two complexes (∼Arc120 and ∼Arc90) connected by short linker DNA (grey)**. The ∼Arc90 complex is bent ∼90^°^ out of plane of the core Arc120 complex.

**Movie M9. EM density flaring of the open-form state, showing that the linker DNA portion is continuous along the length of the Arc207 complex**.

