## Supplementary Material for "Archaeal chromatin ‘slinkies’ are inherently dynamic complexes with deflected DNA wrapping pathways"


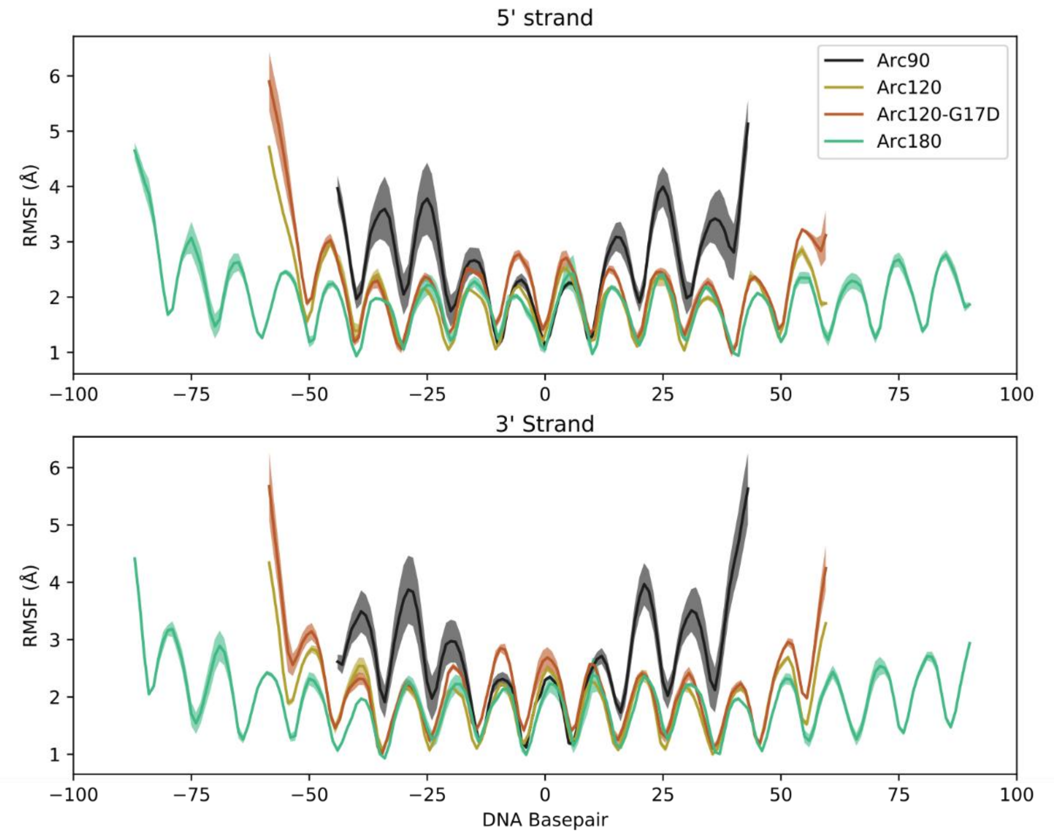


**Figure S1. Root mean-squared fluctuation (RMSF) plots for measured for DNA bases in all four simulated systems.** Average values across the three independent systems are traced by solid lines, and error ranges are calculaged by the standard error of the mean across the three simulations and outlined with shaded regions. RMSF values of the 5' (top) and 3' (bottom) leading strands are shown separately. Periodicity in each plot coincides with DNA turns transitioning between solvent- and protein-facing residues.

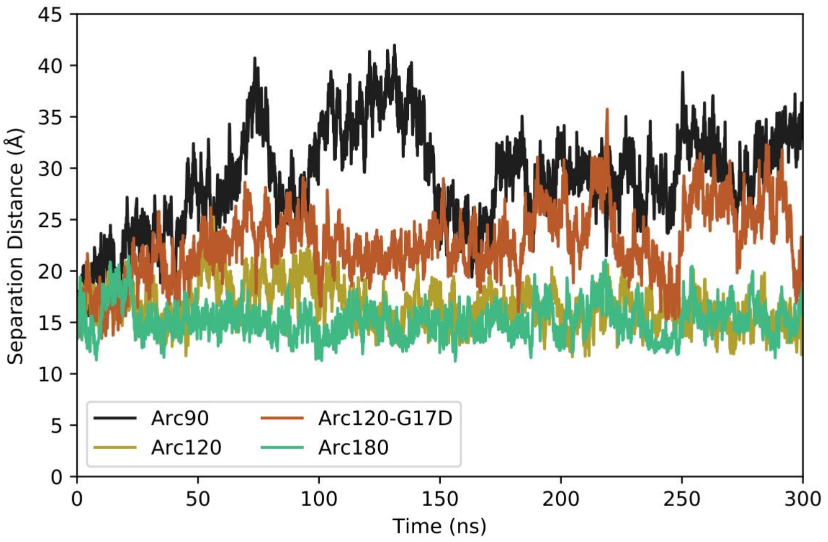


**Figure S2. Representative timeseries for DNA end-to-neighbor separation distances from each system.** The Arc90 system (black) shows both the largest maximum separation, as well as the widest variance in values. The next largest separations and fluctuations are observed in the Arc120-G17D mutant (orange), with the Arc120 wild-type (gold) and Arc180 (teal) systems exhibiting the tightest DNA wrapping.

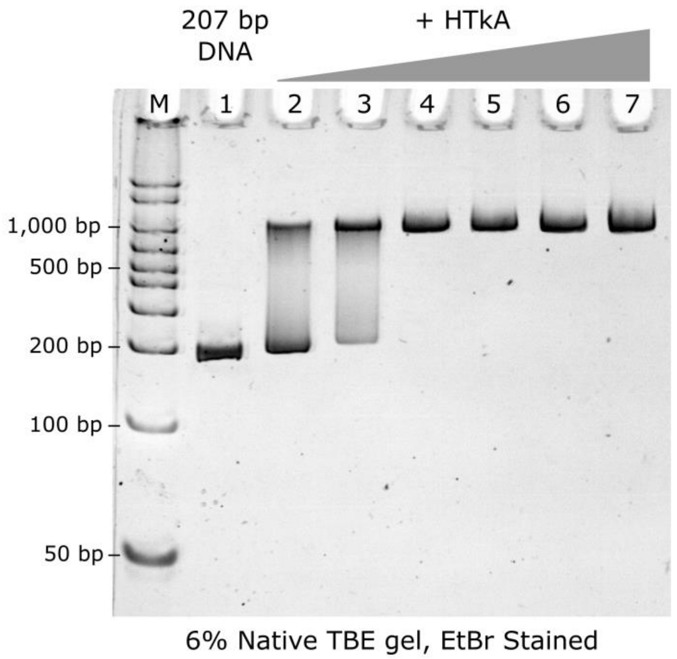


**Figure S3. Titration of HTkA histone on to 207 bp DNA strand, observed by native PAGE gel.** M: Marker; 1: Widom 207 DNA fragment (100 ng); Lanes 2 - 7: 3, 5, 7, 9, 11, and 13 molar equivalents of HTkA histone dimers. Full saturation is observed at the stoichiometric ratio of 7 histone dimers per DNA fragment (lane 4).

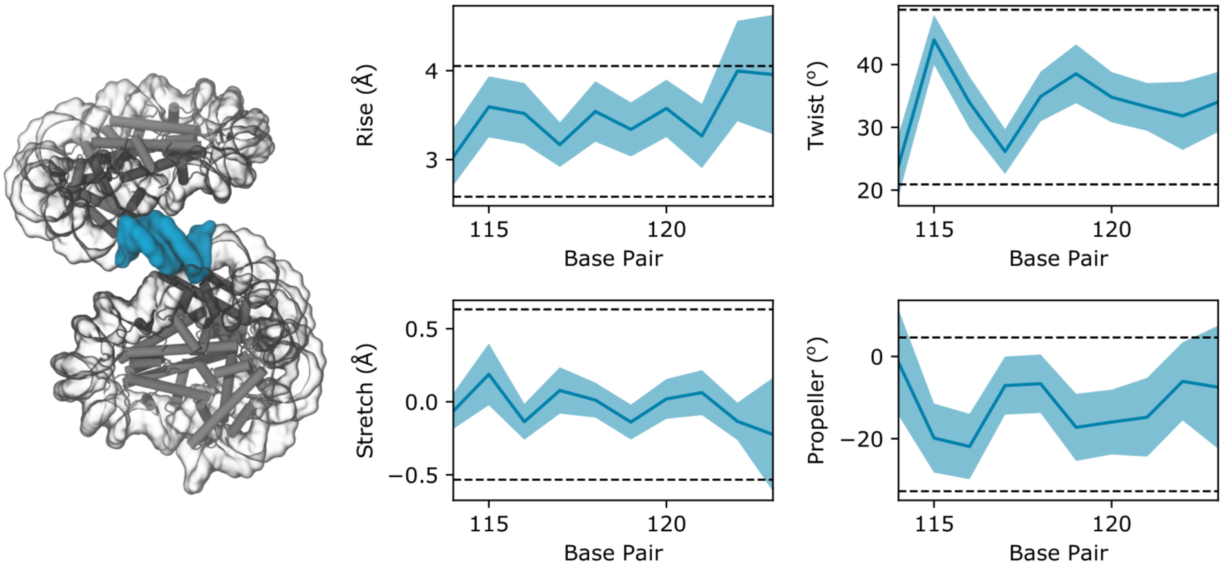


**Figure S4. DNA parameters of the zero-length linker segment calculated from a 10 ns implicit solvent simulation of the Arc207 system restrained to the "open" configuration.** The linker segment is highlighted in blue in the accompanying structure, and **blue** shaded regions in the plot outline the standard deviation of values across the trajectory, whereas the average values are shown by a dark blue line. Horizontal dotted lines define the 2σ range of values observed in the eukaryotic nucleosome (PDB 1aoi).
